# Dynamics of the upper airway microbiome in the pathogenesis of asthma-associated persistent wheeze in preschool children

**DOI:** 10.1101/222190

**Authors:** Shu Mei Teo, Howard HF Tang, Danny Mok, Louise M Judd, Stephen C Watts, Kym Pham, Barbara J. Holt, Merci Kusel, Michael Serralha, Niamh Troy, Yury A Bochkov, Kristine Grindle, Robert F Lemanske, Sebastian L Johnston, James E Gern, Peter D Sly, Patrick G Holt, Kathryn E Holt, Michael Inouye

## Abstract

Repeated cycles of infection-associated lower airway inflammation drives the pathogenesis of persistent wheezing disease in children. Tracking these events across a birth cohort during their first five years, we demonstrate that >80% of infectious events indeed involve viral pathogens, but are accompanied by a shift in the nasopharyngeal microbiome (NPM) towards dominance by a small range of pathogenic bacterial genera. Unexpectedly, this change in NPM frequently precedes the appearance of viral pathogens and acute symptoms. In non-sensitized children these events are associated only with “transient wheeze” that resolves after age three. In contrast, in children developing early allergic sensitization, they are associated with ensuing development of persistent wheeze, which is the hallmark of the asthma phenotype. This suggests underlying pathogenic interactions between allergic sensitization and antibacterial mechanisms.

## INTRODUCTION

Despite advances in modern medicine, acute respiratory tract illnesses (ARIs) continue to be a global health concern. They are a major cause of morbidity and mortality, especially in infants and young children whose immune systems have not yet matured (Ferkol and Schraufnagel, 2014; Zar and Ferkol, 2014), and are the most common reason for antibiotic use in children (Australian Commission on Safety and Quality in Health Care, 2017). The upper airway is a reservoir for microbial communities including viruses, bacteria and fungi, and these have implications for respiratory health and disease. However, current research into the aetiology of ARIs focuses primarily on viruses, most notably respiratory syncytial virus (RSV) and human rhinoviruses (RV). The bacterial microbiome is increasingly recognised as playing an important role in the susceptibility and severity of ARIs, as well as non-communicable respiratory diseases such as asthma (de Steenhuijsen Piters et al., 2015; Durack et al., 2016; Man et al., 2017; Vissers et al., 2014). Infancy is a critical time when microbial colonization may influence an individual’s future respiratory health or disease; indeed, epidemiological data show that repeated ARIs during early childhood are a major risk factor for wheeze and asthma that persist into adulthood (Holt and Sly, 2012).

In recent years, we and others have described the nasopharyngeal microbiome (NPM) in early life (from birth to one or two years of age) (Biesbroek et al., 2014; Bisgaard et al., 2007; Bogaert et al., 2011; Bosch et al., 2017; Teo et al., 2015; Tsai et al., 2015). These independent studies in different human populations have reported strikingly similar findings. Firstly, the NPM appears to be simple in structure, with distinct profiles dominated by a single bacterial operational taxonomic unit (OTU) or genus. A *Staphylococcus*-dominated profile can be observed in early infancy (from one week) but its prevalence decreases sharply over the first year, to be replaced by *Corynebacterium*, *Alloiococcus* (*Dolosigranulum*) or *Moraxella*-dominated profiles, with transient incursions of *Streptococcus* or *Haemophilus*-dominated profiles during ARIs. Secondly, NPM composition influences both microbiota stability and ARI risk and severity. One study of 60 infants from Netherlands in the first two years of life reported a Moraxella-dominated or mixed *Corynebacterium/Dolosigranulum* profile at 1.5 months of age was associated with high NPM stability and low frequency of parent reported ARIs in the subsequent period (Biesbroek et al., 2014). Our study of 234 infants from Australia in the first year of life also found that Moraxella-dominated and Alloiococcus-dominated profiles were more stable than others, but a Moraxella-dominated NPM at two months of age was associated with earlier onset of first ARI (Teo et al., 2015). There were also common environmental correlates of the NPM which differed slightly across studies but included mode of delivery, infant feeding, season, crowding or exposure to other children, recent antibiotic use, and prior infections. Many of these studies, however, were limited to samples collected either during illness or during periods of health, and none have yet elucidated the dynamics of the NPM over the entire preschool period.

Understanding patterns of airway microbial colonization and its association with ARIs and subsequent wheeze phenotypes is an important first step towards the potential manipulation of the microbiome in treating or preventing acute or chronic respiratory disease. In this study, we performed a comprehensive characterization of the largest longitudinal collection of nasopharyngeal samples reported to date – over 3,000 samples from 244 children collected during periods of respiratory health and acute illness over the first five years of life, as part of the prospective Childhood Asthma Study (CAS) (Kusel et al., 2008; Kusel et al., 2006; Kusel et al., 2007; Kusel et al., 2012; Teo et al., 2015). We have previously reported an association between viral-associated lower respiratory infections (LRIs) in infancy, especially those accompanied by fever, and the development of persistent wheeze and asthma in later childhood at five and ten years of age in this cohort, particularly for infants who developed allergic sensitization by age two (Kusel et al., 2007; Kusel et al., 2012). In addition, we recently reported patterns of bacterial colonization in samples collected during the first year of life, and found that specific NPM profiles were associated with ARI symptom severity, independent of the effect of common respiratory viruses (Teo et al., 2015). Here, we address several major research gaps: (i) we examine the relationships and longitudinal dynamics of NPM colonization within children, in health and during ARI episodes, across the first five years of life; (ii) we investigate changes in the association of ARI symptoms with specific NPM taxa over the preschool years; (iii) we address whether the appearance of viral pathogens in the NPM is a harbinger of change in local bacterial populations, or vice versa; and (iv) we investigate the relationship between NPM colonization, early allergic sensitization and future persistent wheeze at five years of age.

## RESULTS

We characterized the bacterial microbiome of 3,014 nasopharyngeal samples from 244 infants in their first five years of life, using 16S rRNA V4 region amplicon sequencing (**Methods**). This included 1,018 “healthy” samples (sampled regularly at 6- or 12-month intervals in the absence of symptoms of respiratory illness; median 5 samples per child, IQR 3-6), 964 samples from upper respiratory illnesses (URIs; median 4 samples per child, IQR 2-6), and 1,032 samples from lower respiratory illnesses (LRIs; median 4 samples per child, IQR 2-7) (**Methods**).

### Composition of upper airway microbiota in the first five years of life

Across all 3,014 samples, the dominant bacterial genera were *Moraxella* (40.1%), *Streptococcus* (13.3%), *Corynebacterium* (12.1%), *Alloiococcus* (11.1%), *Haemophilus* (8.6%), and *Staphylococcus* (4.2%). These made up 89% of all reads, consistent with previously reported results in this cohort for the first year of life (Teo et al., 2015) (Figure 1A, S1A). Each of the six major genera was comprised of multiple operational taxonomic units (OTUs), although the majority were extremely rare. OTU distributions within *Moraxella, Alloiococcus* and *Corynebacterium* were less diverse, with ≤5 OTUs making up ≥97% of all reads from each respective genus at any time period (**Figure S1B**). OTU distributions within *Streptococcus*, *Haemophilus*, and *Staphylococcus* were more diverse, although still dominated by one or two OTUs (**Figure S1B**). The phylogenetic relationships between OTUs are indicated by the tree in Figure 1A, and details of their distribution and predicted species associations are discussed in **Supplementary Text, Figure S2** and **Table S2**.

**Figure 1:**
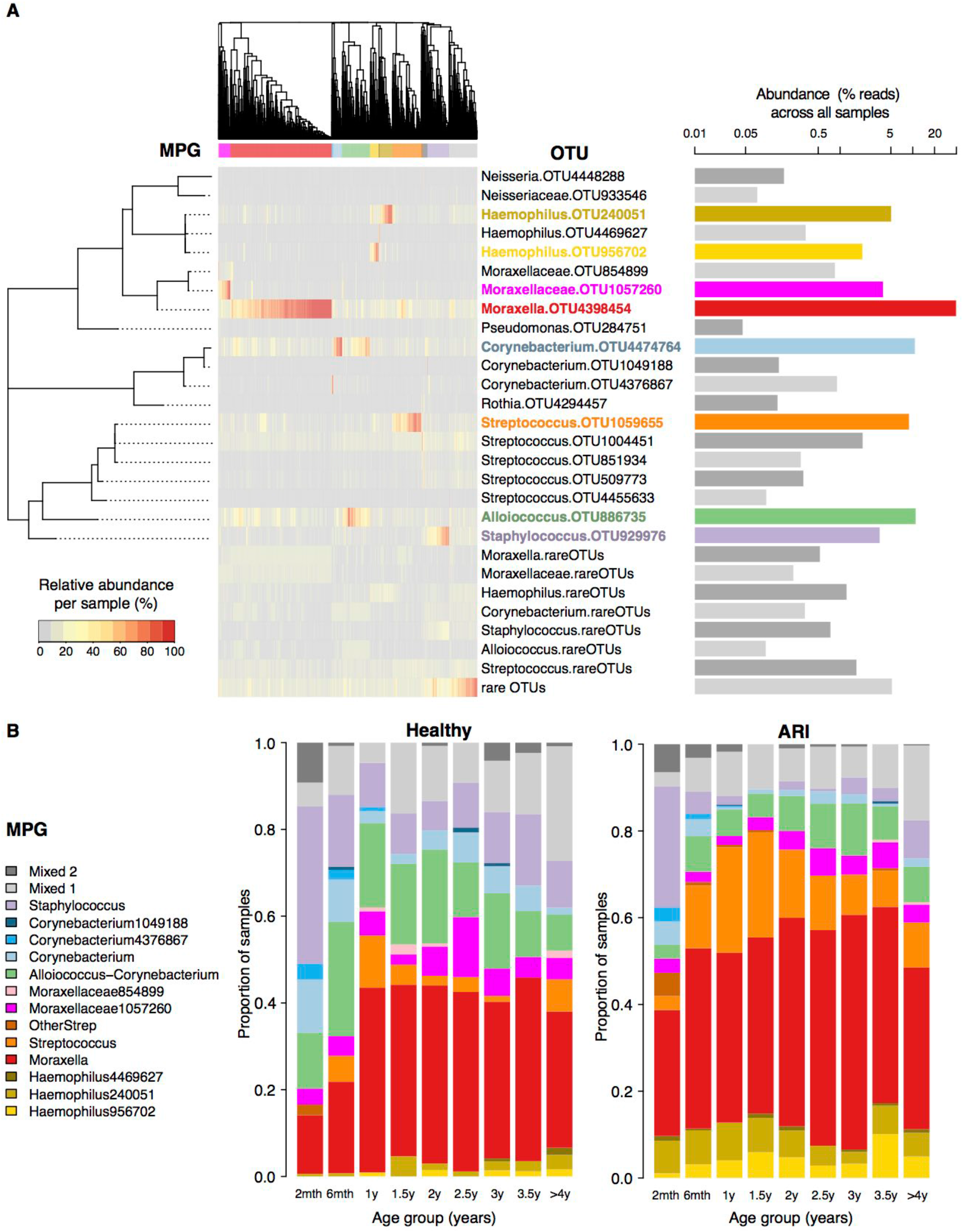
Definition and distribution of microbiome profile groups (MPGs). (**A**) Heatmap shows relative abundances of 21 common operational taxonomic units (OTUs); aggregated values for other OTUs from the common genera; and aggregated values for all other rare OTUs within each sample. Tree to the left shows phylogenetic relationships between the 21 common OTU sequences. Dendrogram at the top indicates complete linkage clustering of Bray-Curtis distances between samples; coloured bars indicate assignment to MPGs based on this clustering (number of MPGs was chosen to maximize the median silhouette value). Barplot to the right shows the total abundance of each OTU or group of OTUs within the whole dataset; OTUs that dominate a common MPG are coloured to match that MPG. (**B**) Distribution of MPGs within each time period, shown separately for healthy and ARI (acute respiratory illness) samples.

The distribution of the relative abundances of common OTUs across samples was highly structured (heatmap in Figure 1A). We used hierarchical clustering to assign each sample to one of 15 microbiome profile groups (MPGs) (Figure 1). Most MPGs were dominated by one OTU, which was used to label each MPG (Figure 1, **Table S1**). The exceptions were two ‘mixed’ MPGs. ‘Mixed1’ MPG contains samples (n=327) in which the common OTUs were all at low abundance, and the vast majority of mixed1 samples (97%) were not dominated by any OTU; the rest were mostly ARI samples dominated by genera that likely represent known respiratory pathogens (*Mycoplasma*, *Bordatella*, *Neisseria*, *Pseudomonas*, *Prevotella*). ‘Mixed2’ MPG (n=51) represents a heterogeneous cluster with no distinct profile. The distribution of MPGs across ages in healthy samples and during ARI is shown in Figure 1B.

Within-sample (alpha) diversity of the NPM increased with age in both healthy and ARI samples, with a noticeable increase after two years of age (GEE linear regression of Shannon’s diversity index vs. age, adjusted for symptom status: p = 0.8 before two years, p = 4x10^−15^ after two years; see Figure 2A). The increasing diversity after two years of age was due to both increasing number and increasing equitability (evenness) of the OTUs (**Figure S3A-B**), and this trend was observed within all MPGs (**Figure S3C**).

**Figure 2:**
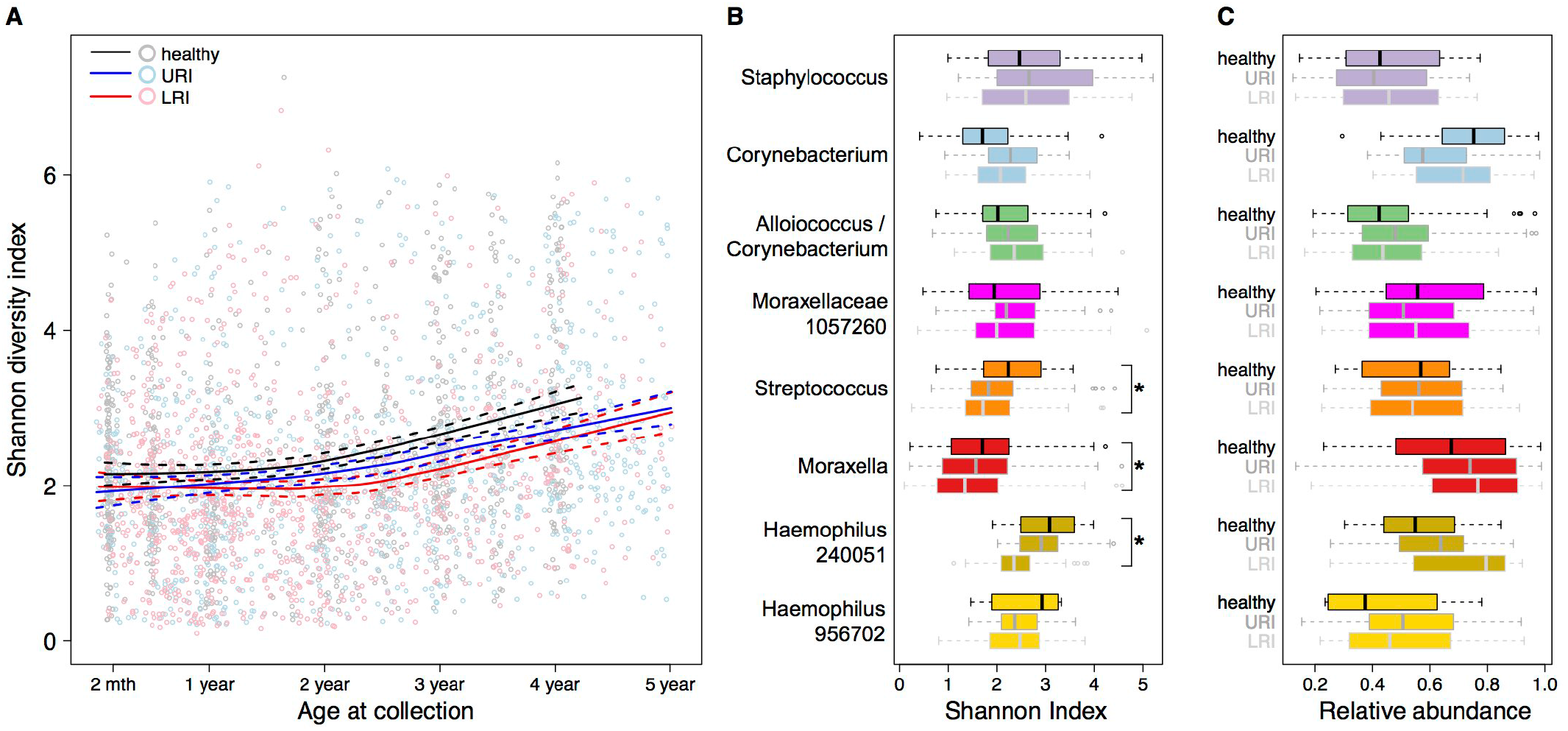
Within-sample diversity is associated with age and acute respiratory illness symptoms. (**A**) Shannon diversity index (SDI) per sample over time, coloured by symptom status as indicated (URI=upper respiratory illness; LRI=lower respiratory illness). Solid lines, loess smoothed curves; dashed lines, 95% confidence intervals. (**B**) SDI distributions within common MPGs. *=FDR adjusted p-value <0.05 in GEE linear regression of SDI against healthy vs. LRI, adjusted for age at collection. (**C**) Relative abundances of the dominant OTU within each MPG (as specified in **Table S1**).

### Upper airway bacteria and respiratory infection

Consistent with our previously reported observations from the first year of life (Teo et al., 2015), ARI was positively associated with MPGs dominated by *Haemophilus* (OR 4.6, p = 1.9x10^−12^), *Streptococcus* (OR 3.9, p = 1.7x10^−17^) or *Moraxella* (OR 1.3, p = 1.8x10^−4^) (adjusting for age, gender, season and recent antibiotics; see **Table S1**). Within these MPGs, the relative abundance of the dominant OTU increased, and the alpha diversity decreased, with symptom severity (comparing healthy, to URI, to LRI; Figure 2B, **Table S3**). Thus overgrowth of these illness-associated taxa accompanies spread of infection to the lower airways, although the direction of causation is not resolvable here.

At the OTU level, the relative abundances of 236 common OTUs (present in >10 %of samples) were significantly different between ARI and healthy samples (absolute difference >1.5-fold and false-discovery rate (FDR) adjusted p-value <0.05). However, these associations were age-dependent and appeared to shift following the increase in within-sample diversity observed from two years of age. Comparing the time periods before year two and on or after the second birthday, a total of 310 OTUs were found to be significantly associated with ARI in at least one interval (absolute difference >1.5-fold and FDR adjusted p-value <0.025; see **Figure S4A**). The majority of *Moraxella, Haemophilus* and *Streptococcus* OTUs were consistently positively associated with ARI in both time periods. *Staphylococcus*, *Corynebacterium* and *Alloiococcus* OTUs were negatively associated with ARI in the first 2 years, but these associations waned after 2 years, particularly for *Corynebacterium* and *Alloiococcus* (**Figure S4A**). Interestingly, we found two *Streptococcus* OTUs that were negatively associated with ARI (OTUs 4365744 and 509773, which show closest match to species *gordonnii* and *thermophilus* / *salivarius* / *vestibularis*, respectively; see **Table S2, Figure S2A**). These were positively correlated with one other, but negatively correlated with *Streptococcus* OTU 1059655, which has closest match to *pneumoniae* / *pseudopneumoniae* (**Figure S5**). The set of five MPGs dominated by ARI-associated *Moraxella*, *Streptoccocus* or *Haemophilus* OTUs are collectively termed here “illness-associated MPGs” (**Table S1**). Amongst the rare OTUs, eight including from genera *Nitriliruptor* and *Bacilllus* were consistently health-associated and often co-occurred together (at median relative abundance 9%) in samples belonging to the *Staphylococcus* MPG, whilst a further eight OTUs including from genera *Porphyromonas*, *Candidatus Aquiluna*, *Clavibacter*, *Mycobacterium, Granulicatella* and *Fusobacterium* were consistently ARI-associated but generally of low abundance across all MPGs (**Figure S4B-C**).

To more precisely estimate how these associations change with age, we performed time varying analysis for eight characteristic OTUs using smoothing splines ANOVA (see Figure 3 and **Methods**). The results showed that the strength of association of *Corynebacterium, Alloiococus* and *Staphylococcus* with healthy samples was greatest in the first 1–2 years; interestingly the association waned towards the null by three years of age for *Corynebacterium* and four years of age for *Staphylococcus*, but *Alloiococcus* changed direction and was significantly associated with ARI in the interval 2.8–3.9 years (mean difference 1.8-fold, p = 0.01) (Figure 3). *Alloiococcus otitidis* in the ear canal has been implicated in otitis media (OM) (Harimaya et al., 2006; Tano et al., 2008), along with *S. pneumoniae* and *H. influenzae* (Ngo et al., 2016). In this cohort, 11% of ARI episodes co-occurred with OM. *Streptococcus* OTU 1059655 abundance in the NPM during ARI was positively associated with concurrent OM from 18 months (mean difference 2.4-fold, p=0.01), but *Alloiococcus* and *Haemophilus* showed no significant associations with OM. The *Alloiococcus–illness* association is further addressed below.

**Figure 3:**
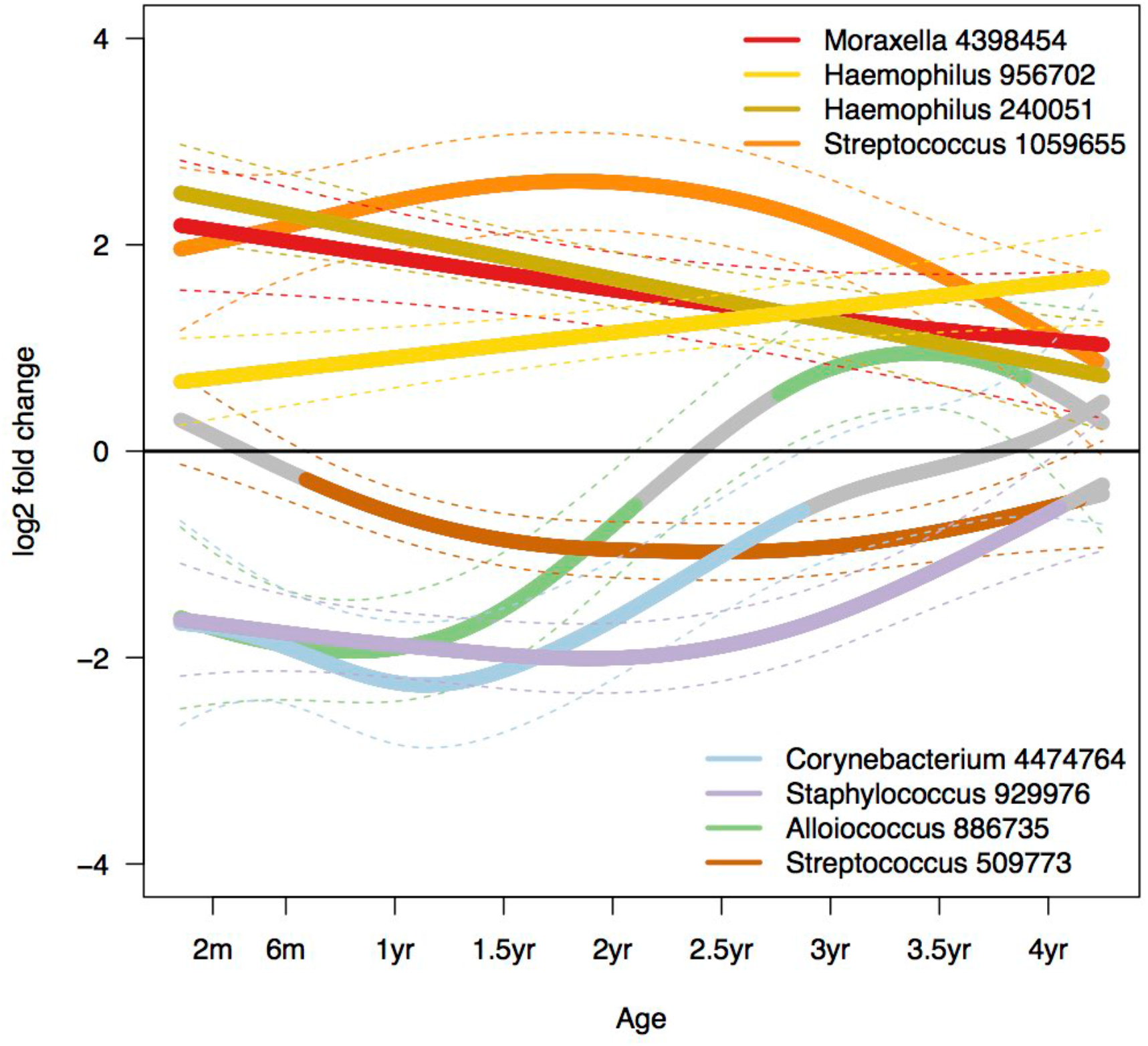
Time varying associations of bacterial taxa with acute respiratory illness symptoms. Log2 fold change (solid lines) and 95% confidence intervals (dashed lines) comparing symptomatic vs. healthy samples, estimated using smoothing splines ANOVA, adjusted for age, season, gender, and any recent antibiotics. Non-significant segments are coloured grey.

### Interactions between bacteria and viruses in the upper airways

We calculated pairwise correlation networks between common OTUs (those present in >10% of samples) using an accelerated FastSpar implementation of the SparCC algorithm (Friedman and Alm, 2012) (see **Methods**). Correlations were calculated separately for samples collected before two years of age and on or after the second birthday; values for the eight most common OTUs are shown in Figure 4A. Illness-associated OTUs of *Moraxella*, *Haemophilus*, and *Streptococcus* formed a group that were all positively correlated with one another in both time periods, and negatively correlated with the health-associated *Streptococcus* OTU 509773 and *Staphylococcus* (Figure 4A). *Corynebacterium* and *Alloiococcus* were strongly correlated with one another in both periods (SparCC correlation=0.68, p = 0.001). Surprisingly, the relationship between these OTUs and those in the illness-associated group (Figure 4A) was complex and changed over time, becoming significantly more positively correlated with age (Figure 4B). *Corynebacterium* and *Alloiococcus* were positively correlated with *Moraxella* throughout, with correlation strength increasing significantly over time (Figure 4A; Fisher’s r to Z transformation comparing *Moraxella–Corynebacterium* and *Moraxella–Alloiococcus* correlations before and after 2 years: p = 9 x 10^−14^ and p = 7 x 10^−16^ respectively). *Corynebacterium* and *Alloiococcus* were negatively correlated with the illness-associated *Haemophilus* and *Streptococcus* OTUs in early years, but became positively correlated (*Streptococcus*) or uncorrelated (*Haemophilus*) in later samples (Figure 4A-B). We hypothesised the increasing co-occurrence with *Moraxella* might explain the increasing association of *Alloiococcus* with ARI symptoms in later years; indeed, adjusting for the abundance of *Moraxella* OTU 4398454 resulted in attenuation of the positive association of *Alloiococcus* with ARI after two years, but not the negative association with ARI prior to two years (**Figure S6**). *Staphylococcus* was positively correlated with the health-associated *Streptococcus* 509773 (SparCC correlation=0.22, p = 0.001) but negatively correlated with the illness-associated group as well as *Corynebacterium* and *Alloiococcus*.

**Figure 4:**
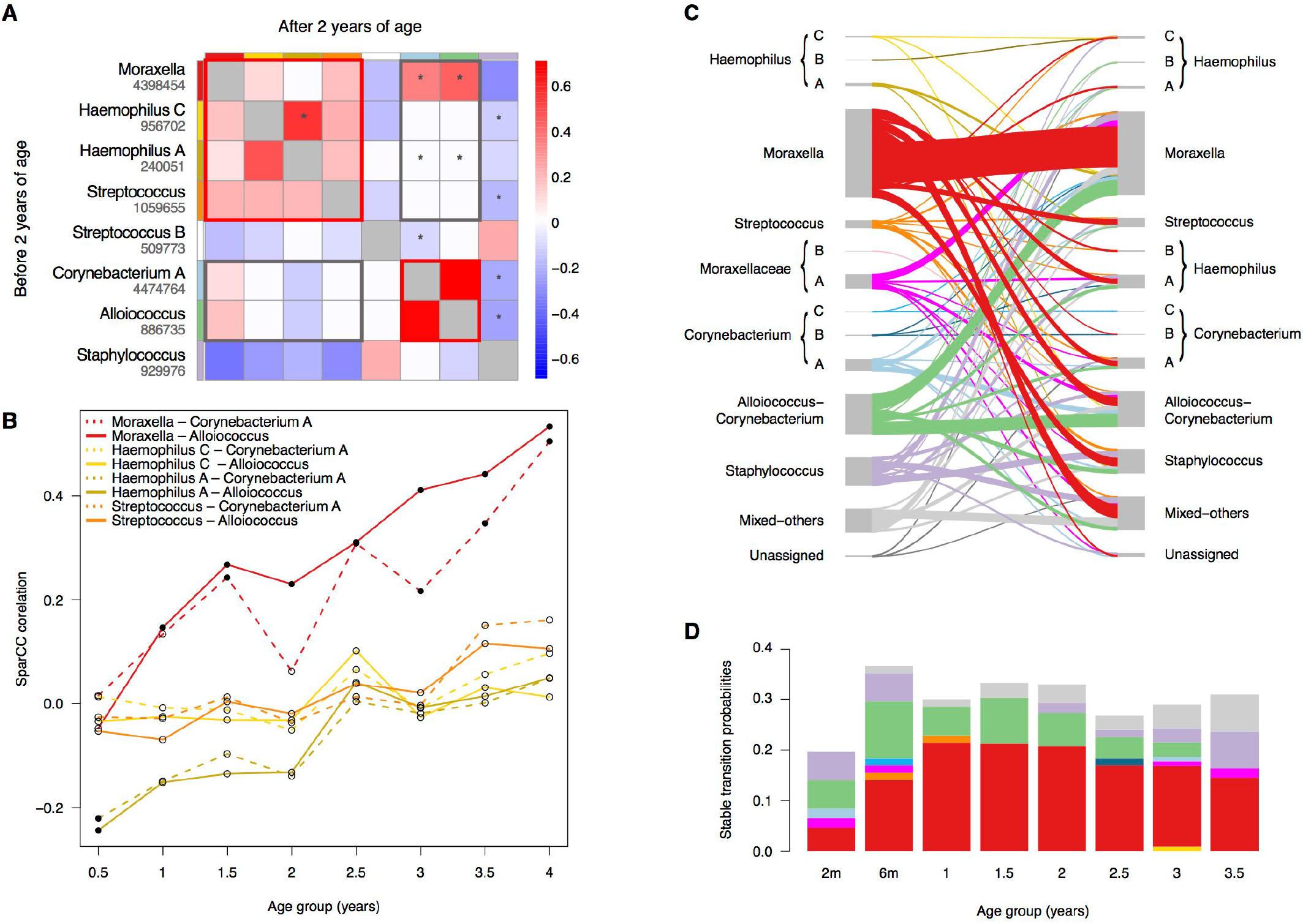
Microbial interaction networks and stability. (**A**) Pairwise correlations among eight characteristic OTUs, estimated using the SparCC algorithm, calculated separately for samples collected up to and including two years of age (lower triangles), and samples collected after two years of age (upper triangles). Cell colours indicate correlation coefficients; non-significant correlations (p>0.001) are coloured white. *Bonferroni-corrected p<0.05/28, testing for change in correlation before and after 2 years of age using Fisher’s z test. (**B**) Correlations between *Alloiococcus* or *Corynebacterium* and *Moraxella* or *Streptococcus* or *Haemophilus* OTUs (bolded black box in A) over half-yearly time periods (Filled circles, significant correlations, p=0.001; empty circles, non-significant correlations, p>0.001). (**C**) Transitions between microbiome profile groups (MPGs) for consecutive pairs of healthy samples collected from the same individuals 6-12 months apart. MPGs are as defined in Figure 1, OTU key: *Haemophilus* A=240051, B=4469627, C=956702; Moraxellaceae A= 1057260, B=854899; *Corynebacterium* A=4474764, B=1049188, C=4376867. (**D**) Proportion of healthy samples collected at each time point, for which the same MPG was detected in the next healthy sample from each individual. Colours indicate the specific MPGs involved, coloured as in panel C.

We tested for common human respiratory viruses in all samples from the first three years of life, using PCR and sequencing (see **Methods**). Viruses were frequently detected in ARI samples (83% and 81% amongst URI and LRI, respectively). Interestingly, the same viruses were also detected amongst 34% of healthy samples, which had been collected after at least one month without ARI symptoms. The presence of virus was significantly associated with illness-associated MPGs (compared to health-associated MPGs) irrespective of symptom status (OR 2.4, p = 1.1x10^−6^ in healthy samples; OR 1.9, p = 1.0x10^−3^ in ARI samples using Fisher’s exact test; see **Figure S7**), suggesting either mutualism, or synergistic effects on symptomatology, between these specific bacterial communities and common respiratory viruses. However, *Streptococcus*, *Moraxella* and *Haemophilus* MPGs were also independently associated with ARI symptoms amongst samples in which no respiratory viruses were detected, or when adjusting for the presence of respiratory viruses or of specific viruses, RSV or RV (Figure 5A-B), suggesting these bacteria contribute to illness in the absence of known respiratory viral triggers.

**Figure 5:**
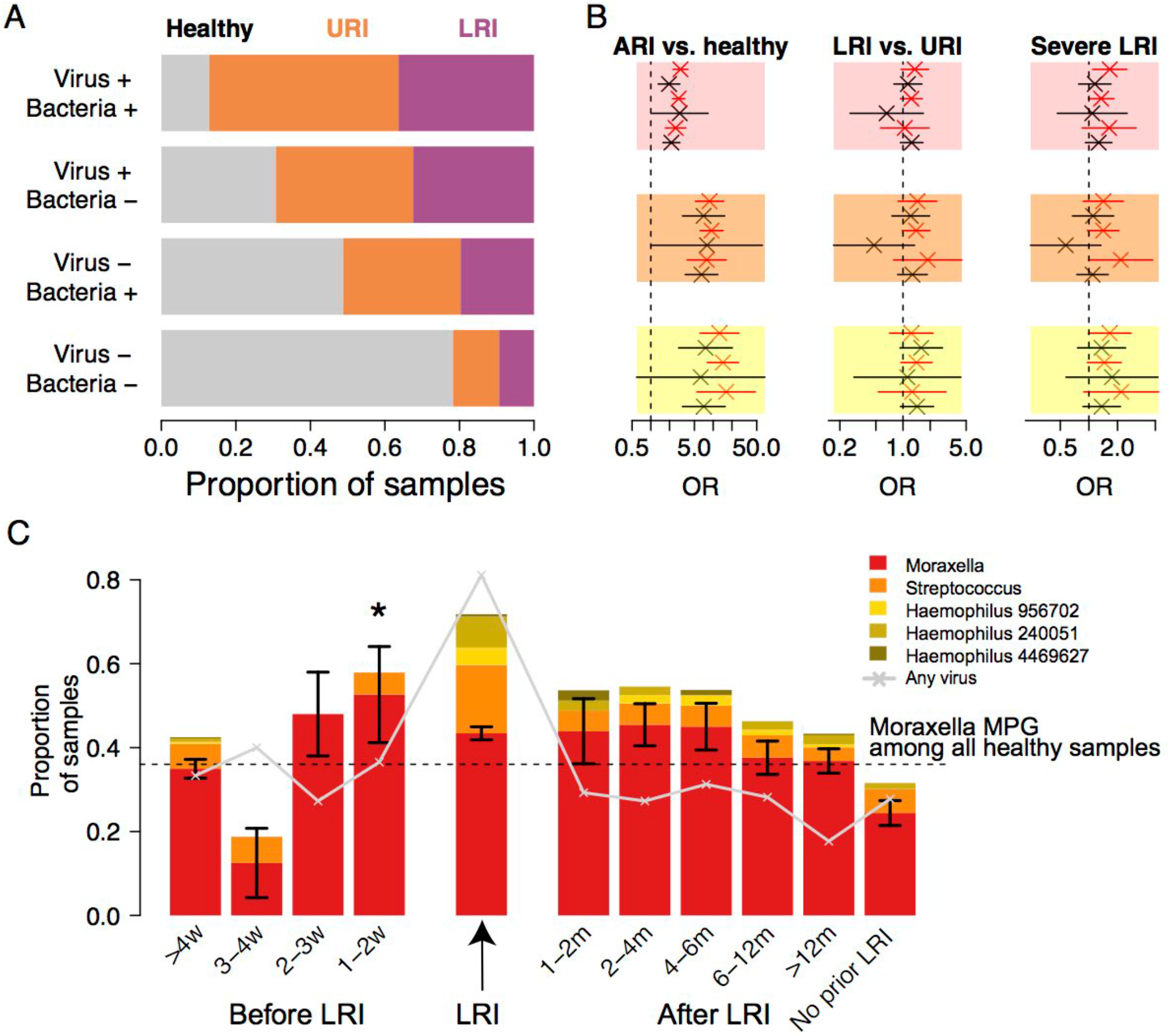
Association of bacteria and viruses with symptoms of acute respiratory illness (ARI). (**A**) Frequency of symptoms (URI=upper respiratory illness, LRI=lower respiratory illness) amongst samples stratified by the presence or absence (+/−) of known respiratory viruses (detected by PCR) and presence or absence (+/−) of bacterial communities assigned to *Moraxella, Streptococcus* or *Haemophilus* microbiome profile groups (MPGs). (**B**) Association of ARI symptoms with specific MPGs, stratified by the presence or absence (+/−) of common respiratory viruses (RV=rhinovirus, RSV=respiratory syncitial virus, Vir=any virus). Odds ratios (OR) and 95% confidence intervals were estimated using generalized estimating equations (GEE) with unstructured correlation and robust standard errors, adjusting for age, gender and season. (**C**) Proportion of healthy samples assigned to *Moraxella*, *Streptococcus* or *Haemophilus* MPGs, stratified by time relative to a recorded LRI episode. Standard error bars are given for the *Moraxella* MPG. We regressed assignment to *Moraxella* MPG against time to LRI (separate models for each time category versus all other healthy samples), and adjusted for time post-LRI, gender, age, season and recent antibiotics; this showed healthy samples collected 1-2 weeks before an LRI were significantly enriched for *Moraxella* MPGs (*p<0.05).

### Stability of the upper airway microbiota within individuals

The NPM is a complex ecosystem that is inherently dynamic as it is continually being shaped by multiple factors, including responses to environmental perturbation and disease status of the host. We therefore examined the effects of external factors, including ARI and antibiotic exposure, on intra-individual NPM dynamics.

We first considered consecutive healthy samples from each individual, excluding sample pairs collected more than one year apart. Overall, the probability of the next consecutive healthy sample sharing the same (non-mixed) MPG as the current sample (i.e. a stable transition) was greater than expected by chance (31% vs 18% for samples collected <6 months apart, binomial test p = 2.3 x 10^−7^; 23% versus 18% for samples collected 6-12 months apart, p = 0.011), indicating some degree of stability of the microbial communities within individuals over time. This stable transition probability was highest for the *Moraxella* (45%) and *Alloiococcus-Corynebacterium* (32%) MPGs, which were the most common states for healthy NPM samples (Figure 4C). Where a sample was assigned to the mixed1 MPG, the probability of the next sample also being designated mixed1 was high (30%). However, the composition of such samples can vary widely, and Bray-Curtis distances between consecutive mixed1 MPG samples were close to the distances between distinct MPGs (**Figure S8**), indicating that the majority of consecutive pairs assigned to mixed1 MPG represent significant shifts in NPM composition rather than stable transitions. The probability of a stable transition to the next time point was significantly lower at 2 months of age than at 6, 12 18 or 24-month time points (Figure 4D; 20% vs 31%; Fisher’s exact test p = 0.03), consistent with the observation of a distinct NPM profile at 2 months (Figure 1B). Transition stability declined after 2 years, and an increasing proportion of transitions involved consecutive samples assigned to the mixed1 MPG (Figure 4D), consistent with the observed increase in diversity after age 2 (Figure 2A). The frequency of persistence of the *Staphylococcus* MPG dropped after 6 months and increased again in the fourth year, consistent with prior observations that maternally-transferred *S. aureus* can be detected in infants, but stable colonisation is not established until the pre-school years (Brown et al., 2014; Jimenez-Truque et al., 2012; Schaumburg et al., 2014). Taken together, these results show the NPM is highly variable in early childhood. Stability of health-associated MPGs was significantly disrupted by the occurrence of LRI during the sampling interval, however antibiotic use did not significantly alter the probability of stable transitions (Table 1).

**Table 1:** Association between MPG instability and intervening illness and antibiotics. We considered pairs of consecutive healthy samples from each individual (excluding the 2-month time point and sample pairs collected >1 year apart), and classified stable transitions as pairs in which the same (non-mixed) MPG present at the first time point was also present at the next healthy sample. Stable transition probabilities for individual MPGs are given in Figure 4D. Here we investigate the impact of extrinsic factors on instability of different groups of MPGs by using a generalized estimating equation (GEE) logistic model with unstructured correlation and robust standard errors, to regress MPG transition instability (stable = 0, unstable = 1) against the presence of intervening ARI, LRI or antibiotics, and further adjusting for age of first sample and difference in ages between samples. Separate models were fit for groups of MPGs associated with acute respiratory illness or respiratory health (as defined in **Table S1**), and for each of the two most common MPGs (*Moraxella* and *Alloiococcus-Corynebacterium*). Values shown are OR (95% CI) and p-values associated with the intervening factor variable (ARI, LRI, antibiotics) in each model; OR >1 indicates increasing instability, OR <1 decreasing instability (i.e. increasing stability).

| Data subset | N | Stable | Instability |  |  |
| --- | --- | --- | --- | --- | --- |
|  |  |  | Any ARI | Any LRI | Any antibiotics |
| All samples | 513 | 31% | 1.3(0.83-2.1),<br>p=0.24 | 1.6(0.95-2.5),<br>p=0.076 | 0.78(0.5-1.2),<br>p=0.3 |
| Illness-associated MPGs | 230 | 41% | 1.3(0.62-2.6),<br>p=0.52 | 0.74(0.41-1.4),<br>p=0.33 | 0.73(0.41-1.3),<br>p=0.3 |
| *Moraxella* MPG | 201 | 45% | 1.2(0.56-2.8),<br>p=0.59 | 0.66(0.34-1.3),<br>p=0.21 | 0.88(0.46-1.7),<br>p=0.68 |
| Health-associated MPGs | 190 | 24% | 1.9(0.9-4.1),<br>p=0.09 | **13(3.2-57),**<br>**p=0.00042** | 0.84(0.32-2.2),<br>p=0.72 |
| *Alloiococcus-Corynebacterium* MPG | 92 | 30% | 1.6(0.56-4.5),<br>p=0.39 | **34(3.8-300),**<br>**p=0.0016** | 0.44(0.11-1.8),<br>p=0.26 |

### Association of upper airway microbiota with lower respiratory illness and subsequent wheeze

We investigated whether LRIs were associated with prior colonization by the illness-associated *Moraxella*, *Streptococcus* and *Haemophilus* MPGs, and how long these MPGs persisted after an incident LRI. Healthy samples collected 1-2 weeks prior to an LRI were not enriched for viruses (~30% frequency of virus detection, vs 34% across all healthy samples and ~80% during LRI), but were significantly enriched for the *Moraxella* MPG (GEE logistic regression of assignment to *Moraxella* MPG on time to LRI, 1-2 weeks compared to all other healthy samples and adjusted for time post-LRI, gender, age, season, recent antibiotics: OR 6.2 [95% CI, 1.4 – 28], p = 0.017; further adjusted for viruses: OR 5.9 [1.3 – 26], p = 0.019) (Figure 5C), as well as *Moraxella* abundance (GEE linear regression of log *Moraxella* OTU abundance, p = 0.025; further adjusted for viruses: p = 0.04) (see **Methods**). There were no significant differences in proportions of *Streptococcus* or *Haemophilus* MPGs nor *Streptococcus* or *Haemophilus* abundance, however these MPGs were rare (~7%) in healthy samples. We were unable to assess short-term changes in the NPM following LRI, as our criteria for healthy sample collection required the absence of ARI symptoms for at least 4 weeks; however, the *Moraxella* MPG exhibited declining frequency with increasing time post-LRI, and remained enriched until six months post-LRI (Figure 5C). Interestingly, there was no evidence of a difference in MPG distribution before or after URI (**Figure S9**).

We have previously shown in this cohort that the risk of chronic wheeze at five years of age is significantly associated with the number of LRIs in the first year of life. This was especially the case for the number of febrile LRIs among children with allergic sensitization by age two (Kusel et al., 2007; Kusel et al., 2012; Teo et al., 2015). Here, we investigated whether presence of the illness-associated *Moraxella*, *Streptococcus* and *Haemophilus* MPGs during the first four years of life was predictive of LRI intensity during the same period, and/or wheeze at age five. For each child, we calculated the combined frequency of these MPGs amongst healthy NPM samples over different time periods (**Methods**). Among children with early allergic sensitization (defined as allergen-specific IgE > 0.35 kU/L by two years of age, see Methods), frequent ARI-associated MPGs (≥50% of healthy NPM samples) during the first two years of life was significantly positively associated with the number of LRIs experienced in the same period (Table 2). Importantly, among these early sensitized children, the frequency of illness-associated

**Table 2:** Associations between proportion of illness-associated MPGs in healthy (asymptomatic) samples and LRI frequency. We modelled the proportion of illness-associated *Moraxella, Haemophilus* and *Streptococcus* MPGs present amongst healthy samples (as a binary variable; ≥50% vs. <50%) on LRI or febrile LRI frequency using logistic regression. Separate models were fit for different time periods, and for children with and without early allergic sensitization by 2 years of age (defined as any allergen-specific IgE > 0.35 kU/L; allergens tested: house dust mite, cat epithelium / dander, peanut, foodmix, couch grass, rye grass, mould mix, infant phadiatop). Values indicate odds ratios (95% CI) and p-values for the association between LRI count (during the years specified by the ‘Yr’ column) and illness-associated MPG frequency ≥50% (amongst healthy samples collected during the period specified in the column header, i.e. 6m–2y or 2.5y–4y).

|  | Yrs | Early allergic sensitization |  | All other children |  |
| --- | --- | --- | --- | --- | --- |
|  |  | MPGs 6m–2y<br>(N=74) | MPGs 2.5y–4y<br>(N=83) | MPGs 6m–2y<br>(N=65) | MPGs 2.5y–4y<br>(N=65) |
| # LRI | 0-1 | 1.6(1-2.6),<br>p = 0.043 | 1.1(0.83-1.5),<br>p = 0.49 | 1.5(0.95-2.3),<br>p = 0.081 | 1.1(0.79-1.5),<br>p = 0.57 |
|  | 1-2 | 1.5(1-2.2),<br>p = 0.036 | 1.1(0.81-1.5),<br>p = 0.6 | 1.4(0.91-2.1),<br>p = 0.12 | 1.2(0.81-1.7),<br>p = 0.38 |
|  | 2-4 | 1.4(1-1.9),<br>p = 0.032 | 1.1(0.79-1.4),<br>p = 0.72 | 0.99(0.79-1.2),<br>p = 0.93 | 1.1(0.87-1.3),<br>p = 0.47 |
| # Febrile LRI | 0-1 | 2.5(1-6.3),<br>p = 0.049 | 1.2(0.63-2.4),<br>p = 0.53 | 2.2(0.77-6.1),<br>p = 0.14 | 0.84(0.36-2),<br>p = 0.69 |
|  | 1-2 | 1.8(0.85-3.6),<br>p = 0.13 | 0.59(0.3-1.2),<br>p = 0.13 | 2.4(0.65-8.9),<br>p = 0.19 | 1.8(0.71-4.6),<br>p = 0.21 |
|  | 2-4 | 1.8(0.88-3.9),<br>p = 0.11 | 0.99(0.5-2),<br>p = 0.98 | 1.6(0.74-3.5),<br>p = 0.23 | 1.4(0.76-2.5),<br>p = 0.3 |

MPGs in the first two years of life was independently associated with chronic wheeze at five years of age (Figure 6A), even after adjusting for LRI frequency and type (Table 3). Notably, among non-sensitized children, the frequency of illness-associated MPGs was not associated with chronic wheeze at five years but was significantly positively associated with the transient wheeze phenotype (defined as any wheeze in the first three years of life but no wheeze in the fifth year) (Figure 6, Table 3).

**Figure 6:**
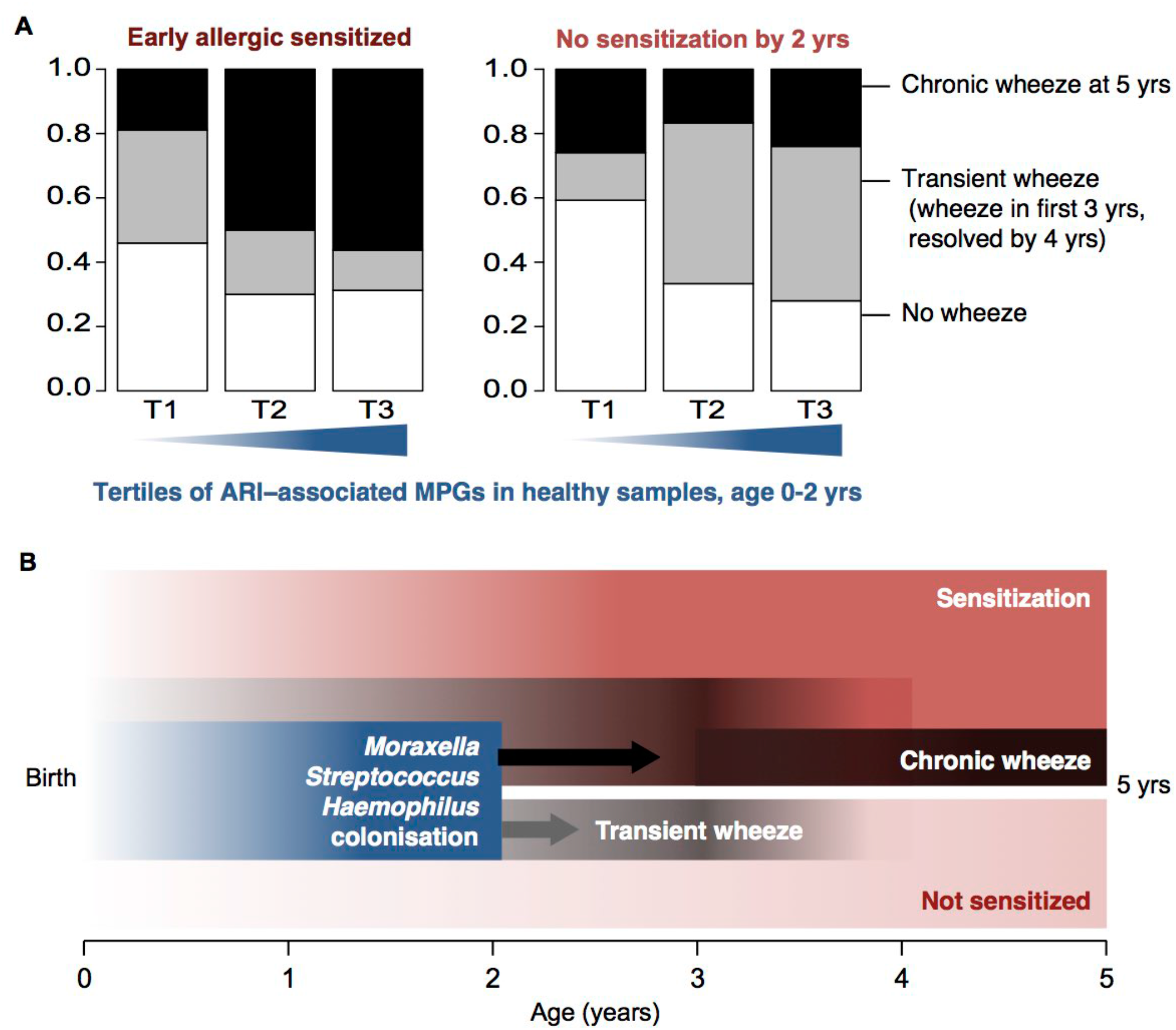
NPM associations with wheeze. (**A**) Frequency of pre-school wheeze phenotypes (y-axis), stratified by frequency of *Moraxella*, *Streptococcus* or *Haemophilus* MPGs amongst healthy samples collected from 6 months to 2 years of age (x-axis, in quartiles). Data are shown separately for 73 children who were allergic sensitized by 2 years of age, and 64 who were not. (**B**) Schematic diagram illustrating that among early sensitized children, the intensity of asymptomatic colonization by *Moraxella, Streptococcus* or *Haemophilus* in the first two years of life was associated with increased risk of chronic wheeze at age five; however, in children who were not sensitized early, it was associated only with transient early wheeze.

**Table 3:** Prediction of subsequent wheeze phenotypes based on proportion of illness-associated MPGs amongst healthy samples during the first two years of life. Logistic regression of wheeze phenotype (Y) against tertiles of the proportion of illness-associated *Moraxella, Haemophilus* and *Streptococcus* MPGs (%MPG), adjusting for illness frequency (X). Separate models were fit for children with and without early allergic sensitization by 2 years of age (defined as any allergen-specific IgE > 0.35 kU/L; allergens tested: house dust mite, cat epithelium / dander, peanut, foodmix, couch grass, rye grass, mould mix, infant phadiatop).

| Data subset | Y | Model | Estimated OR (95% CI) and p-value |  |
| --- | --- | --- | --- | --- |
|  |  | X | % MPG | X |
| Early sensitized children (N = 73) | Wheeze at 5y | None | **2.5 (1.3-4.6), p = 0.0054** | NA |
|  |  | # LRI (yr 1) | **2.2 (1.1-4.2), p = 0.018** | 1.5 (0.94-2.5), p = 0.085 |
|  |  | # LRI (yr 2) | **2.3 (1.2-4.3), p = 0.013** | 1.3 (0.89-1.8), p = 0.18 |
|  |  | # LRI (yr 3-4) | 2 (0.94-4.2), p = 0.073 | **2.4 (1.5-3.9), p = 0.00059** |
|  |  | # febrile LRI (yr 1) | **2.1 (1.1-4.1), p = 0.026** | **2.9 (1.1-7.5), p = 0.032** |
|  | Transient wheeze | None | 0.5 (0.23-1.1), p = 0.074 | NA |
|  |  | # LRI (yr 1) | **0.44 (0.19-0.99), p = 0.047** | 1.4 (0.8-2.3), p = 0.26 |
|  |  | # LRI (yr 2) | **0.41 (0.17-0.96), p = 0.039** | 1.3 (0.93-1.9), p = 0.12 |
|  |  | # LRI (yr 3-4) | 0.54 (0.25-1.2), p = 0.12 | 0.86 (0.6-1.2), p = 0.39 |
|  |  | # febrile LRI (yr 1) | 0.48 (0.22-1.1), p = 0.068 | 1.3 (0.45-3.5), p = 0.67 |
| All other children (N = 64) | Wheeze at 5y | None | 0.94 (0.5-1.8), p = 0.86 | NA |
|  |  | # LRI (yr 1) | 0.93 (0.49-1.8), p = 0.83 | 1.1 (0.73-1.5), p = 0.74 |
|  |  | # LRI (yr 2) | 0.91 (0.46-1.8), p = 0.78 | 1.5 (0.99-2.2), p = 0.059 |
|  |  | # LRI (yr 3-4) | 1.2 (0.49-3), p = 0.66 | **3.8 (1.8-8.3), p = 0.00062** |
|  |  | # febrile LRI (yr 1) | 0.94 (0.49-1.8), p = 0.84 | 1.2 (0.42-3.2), p = 0.76 |
|  | Transient wheeze | None | **2.2 (1.2-4.1), p = 0.014** | NA |
|  |  | # LRI (yr 1) | **2.2 (1.1-4.2), p = 0.022** | **1.5 (1-2.2), p = 0.027** |
|  |  | # LRI (yr 2) | **2.2 (1.2-4.1), p = 0.014** | 1.1 (0.75-1.6), p = 0.62 |
|  |  | # LRI (yr 3-4) | **2.3 (1.2-4.6), p = 0.014** | **0.53 (0.3-0.92), p = 0.024** |
|  |  | # febrile LRI (yr 1) | **2.2 (1.1-4.3), p = 0.018** | **3.4 (1.2-9.5), p = 0.018** |

## DISCUSSION

This study presents the first comprehensive, longitudinal characterization of the upper airway microbiome in a cohort followed from birth to five years of age, and its association with episodes of ARI, allergic sensitization and subsequent wheezing phenotypes. We found the NPM from birth to five years of age remains dominated by six common genera (Figure 1) and has yet to converge to an adult-like NPM, which is characterised by much greater alpha diversity, lack of *Moraxella* and *Corynebacterium*, and much lower biomass (Stearns et al., 2015). This is in contrast to the oropharynx (Stearns et al., 2015), or the gut microbiome, which matures to an adult-like state by three years of age (Yatsunenko et al., 2012). Consistent with previous observations (Biesbroek et al., 2014), we found a constant level of NPM diversity over the first two years of life, followed by a period of increasing diversity – in terms of both number and equitability of OTUs – for at least three years (**Figures 2A, S2**). This increasing diversity coincided with a change in the relationship between the NPM and respiratory disease, whereby negative associations between MPGs and ARI became attenuated (as in the case of *Corynebacterium*) or changed direction to become positively associated with ARI (*Alloiococcus*) (Figure 3). In the latter case, this appears to be driven by an increasing alliance with *Moraxella* (**Figures 4, S6**), which itself was ARI-associated. *Moraxella* establishes biofilms that enhance the co-survival of pathogens such as *Streptococcus pneumoniae* and *Haemophilus influenzae* (Pearson et al., 2006; Perez et al., 2014). It is not yet clear whether the negative associations of certain taxa with ARI denote active protective effects, or simply the lack of pathogenic drivers of symptoms; however there is some evidence from murine models that pre-exposure to *Corynebacterium* can provide some resistance against RSV infection (Kanmani et al., 2017).

Our longitudinal data show the NPM can be highly dynamic within individuals. However there was some stability even between samples collected 6 or 12 months apart (Figure 4C-D), especially for the MPGs dominated by *Moraxella* or *Alloiococcus* and *Corynebacterium*, which appear to be stable colonizers of the nasopharynx of children. Notably, stability of the *Alloiococcus–Corynebacterium* MPG was significantly reduced by LRI episodes, which are typically associated with an influx and/or overgrowth of *Moraxella, Streptococcus* or *Haemophilus* that can presumably destabilise the bacterial community. This is consistent with a recent study that reported reduced stability of the NPM during infancy among children who experienced more than two ARIs in the first year of life (Bosch et al., 2017). Ultimately, more comprehensive description of natural NPM dynamics, including detailed assessment of resilience to exogenous agents, will require higher resolution sampling (weekly or daily) and would also benefit from larger cohorts.

Throughout the first five years of life, NPM samples collected during ARIs showed a greater abundance of, and were more commonly dominated by, specific *Streptococcus, Moraxella* and *Haemophilus* OTUs (**Figures 1, 3**; **Table S1**), consistent with expectations regarding common respiratory pathogens *S. pneumoniae, M. catarrhalis* and *H. influenzae* to which these OTU sequences were most closely related (**Table S2**). The relative abundances of these OTUs were significantly correlated with one another (Figure 4A); we hypothesise this is related to the protection provided by the *Moraxella* biofilm (Tan et al., 2007), which can release outer membrane surface proteins that protect other bacteria from complement-dependent killing. Other groups have previously reported reduced upper airway microbial diversity during or prior to ARIs (Frank et al., 2010; Santee et al., 2016; Yi et al., 2014); our data supports this, both in terms of enrichment of a small number of community profiles (MPGs) during ARI, and a higher abundance of ARI-associated OTUs and lower alpha diversity within these MPGs compared to that observed in the absence of ARI symptoms (**Figures 2B, S3; Tables S1, S3**). We therefore propose that overgrowth of these particular taxa may tip the balance towards respiratory symptomatology, either by direct action as invasive pathogens or via indirect dysregulation of the local immunological milieu. Such dysregulation may increase the likelihood of a primary viral infection of the nasopharyngeal mucosa, or subsequent spread of infection to the lower airways, as suggested in our earlier study on this cohort during infancy (Teo et al., 2015). This is further supported by the increased prevalence of *Moraxella* in asymptomatic samples collected 1-2 weeks before an LRI (Figure 5C). Most LRIs (>80%) had a known respiratory virus present, and this is likely the trigger for acute symptoms. However, the lack of enrichment for viruses, but enrichment for *Moraxella*, in the 1-2 weeks preceding LRI suggests that having the bacteria present when the virus is encountered increases the likelihood of severe respiratory illness. While our study had insufficient power to detect similar effects for *Streptococcus* and *Haemophilus*, due to low colonization frequency in our cohort, there is a large body of evidence accumulating around specific mechanisms of interaction between human respiratory viruses (mainly RV, RSV and influenza) and *Streptococcus pneumoniae*, *Haemophilus influenzae* and *Moraxella catarrhalis*; including both viral promotion of bacterial colonization and outgrowth (for which we see evidence in the form of increased abundance of pathogenic genera in ARIs), and bacterial promotion of viral receptor expression on host cells (Bosch et al., 2017; Brealey et al., 2015). While the present study cannot address specific mechanisms, it provides evidence for interactions in both directions, and demonstrates that bacterial colonization influences subsequent ARI throughout infancy and early childhood.

Finally, we found a significant relationship between asymptomatic colonization of the upper airways by certain MPGs (*Streptococcus*, *Haemophilus* and *Moraxella*) in the first two years of life and later wheezing phenotypes, conditional on early allergic sensitization (Figure 6, Table 3). In early sensitized children, asymptomatic colonization of the upper airways by illness-associated MPGs increased risk of chronic wheeze at five years of age; however, in children who had not developed early allergic sensitization, it was associated only with transient early wheeze, which resolved by the fourth year of life. Furthermore, in the early sensitized children, the frequency of asymptomatic colonization with illness-associated MPGs was also associated with recurrence of LRIs, particularly those accompanied by fever, throughout the first 4 years of life (Table 2). Notably however, whilst frequency of LRI is associated with five-year chronic wheeze (Kusel et al., 2007; Teo et al., 2015), the effect of bacterial colonization on five-year wheeze remained after adjusting for LRI (Table 3). It has been suggested that the pathogenic bacterial species *S. pneumoniae*, *H. influenzae* and *M. catarrhalis* induce local immunoinflammatory responses in the upper airways of neonates, which in the case of *M. catarrhalis* and *H. influenzae* include upregulation of a mix of Th1/Th2/Th17 cytokines (Folsgaard et al., 2013). However, how these immune responses differ between sensitized and non-sensitized children is incompletely understood. We have suggested, based on previous studies in this and other asthma-risk cohorts (Holt & Sly 2012), that the increased severity of these episodes in children with allergic sensitization is due in part to interactions between infection-associated Type 1 IFN-mediated and allergy-associated Th2-mediated inflammatory pathways which compromise their capacity to efficiently clear respiratory pathogens, thus worsening ensuing airway inflammation and resultant immunopathology. Conversely, host immune defense mechanisms in those who are non-sensitized are not compromised by these interactions, and they accordingly experience only transient illnesses.

In conclusion, this study suggests that the microbiota of the upper airways is an important determinant of the susceptibility, frequency and severity of ARI in early childhood. In conjunction with early allergic sensitization, the dominating presence of illness-associated MPGs (*Streptococcus*, *Haemophilus*, and *Moraxella*) in the upper airways is a significant risk factor for persistent wheeze in school-age children, which is the hallmark of the asthmatic phenotype. This observation is of potential importance in relation to early detection and prevention of asthma. In particular, sensitized children in this cohort already showed elevated levels of allergen-specific IgE production from six months of age (Holt et al., 2010), suggesting a high-risk group could be identified in infancy. Airway microbiome monitoring and potentially modification might be beneficial for this high-risk group, in reducing the risk of lower respiratory infection, the repeated occurrence of which is closely linked to asthma development.

## METHODS

### Sample and data collection

This study is part of the Childhood Asthma Study (CAS) – a prospective community-based cohort of 244 children at high risk of allergic sensitization who were followed prenatally until five years of age with the goal of identifying risk factors for allergic diseases, as previously described (Kusel et al., 2006; Kusel et al., 2007; Kusel et al., 2012; Teo et al., 2015). Healthy nasopharyngeal (NP) samples were collected at planned half-yearly visits (first sample from about 2 months of age, subsequently at 6 months, 1 year, and so on), in the absence of any symptoms of acute respiratory illness (ARI) for at least 4 weeks. In addition, parents were to contact a study clinician at the onset of any ARI symptoms, at which point a study nurse visited the family within 48 hours to collect a NP sample from the child and interview the parents on the symptoms and medications used in relation to the illness. ARIs were classified as a lower respiratory illness (LRI) if accompanied by wheeze or rattly chest; or an upper respiratory illness (URI) otherwise. A total of 1943 healthy samples, 2579 URI samples, and 1056 LRI NP samples were collected and divided into aliquots that were cryofrozen for later analysis. Parents kept a daily record of any medication used, from which antibiotic exposure information was extracted, and completed yearly questionnaires during face-to-face interviews. Blood samples were collected from each child at 6 months, 1, 2, 3, 4 and 5 years of age, and positive sensitization status at each timepoint was defined as serum IgE levels > 0.35 kU/L to house dust mite, cat epithelium and dander, peanut, foodmix, couch grass, rye grass, mould mix, or infant phadiatop (details of which were described in (Holt et al., 2010)).

### Bacterial 16S profiling

One aliquot from each of 1331 healthy, 996 URI and 1055 LRI samples were prepared for bacterial 16S rRNA amplicon sequencing. DNA extraction, amplification of the V4 rRNA region, and amplicon sequencing via Illumina MiSeq were performed as previously described (Teo et al., 2015). Paired end reads were merged using Flash version 1.2.7 (Magoc and Salzberg, 2011) with read length 151 base pairs (bp) and expected fragment length 253 bp. The merged reads were quality filtered as follows: ≤3 low-quality bp (Phred quality score < 3) allowed before truncating a read, ≥189 consecutive high-quality bp, sequences with any N characters were discarded. Reads were clustered into operational taxonomic units (OTUs) using the closed reference OTU picking method in QIIME v1.7 using the Greengenes 99% reference database version 13_05. Mean of 1% of reads per sample had no match to the Greengenes database and were excluded from further analysis (except for alpha diversity calculations, as described below). Negative control samples had a median of >1500 (taxonomy-assigned) reads (interquartile rage 900 – 2300), while NP samples had a median of >147K reads (IQR 45K – 230K). We therefore removed 142 NP samples with <3000 taxonomy-assigned reads, and also 226 healthy samples that did not fulfil the criterion of >4 weeks after an illness episode, leaving 3014 samples (1018 healthy samples and 1996 ARI samples) for further analysis. Because entries in the Greengenes database may be identical in the V4 subregion that we sequenced, it is possible for identical read sequences to be assigned to different Greengenes OTUs. We therefore merged counts for OTUs that were identical in the sequenced V4 region (identified by extracting the sequence between the forward and reverse primer sequences), as shown in **Figure S2**. Read counts were corrected for OTU-specific copy number using Picrust v1.0 (Langille et al., 2013) using the pre-computed copy number estimates for Greengenes OTUs version 13_05; and relative abundances were calculated by normalising to the total taxonomy-assigned reads for each sample. Phylogenetic analyses were conducted by BLAST searching the NCBI 16S rRNA database using representative V4 region sequences from the common taxa (**Figure S1B**) to identify similar sequences. For each genus, the sequences were aligned using Muscle and a maximum-likelihood tree constructed using PhyML (**Figure S2**); these were used to identify the closest known species for each common OTU (**Table S2**).

### Clustering into microbiome profile groups (MPGs)

Samples were assigned to microbiome profile groups (MPGs) based on hierarchical clustering of OTU relative abundances, using Bray-Curtis dissimilarity as the distance metric and complete linkage (implemented in the R function *hclust)*. These analyses included all common OTUs (defined as mean relative abundance >0.1%, present in >20% of samples, and dominating (>50%) at least one sample); aggregated counts of other OTUs from each of the major genera (*Moraxella, Streptococcus, Haemophilus*, *Alloiococcus*, *Corynebacterium*, *Staphylococcus*) and family Moraxellaceae; and a final group consisting of aggregated counts of all other OTUs (labelled ‘rare OTUs’; see rows in Figure 1). The number of clusters (i.e. unique MPGs) was chosen to maximise the median silhouette value. MPGs were named based on the dominant genus or OTU, as indicated in **Table S1**.

### Bacterial diversity analysis

Alpha (within-sample) diversity was assessed using Shannon’s diversity index measure, which takes into account both number and relative abundance of the OTUs. For this analysis we processed the merged and filtered reads in a different manner. First we ran an additional chimera check using UCHIME, then assigned OTUs using the subsampled open reference OTU picking framework in QIIME using the Greengenes 97% OTU reference set (Caporaso et al., 2010). Briefly, the input sequences were prefiltered against the reference set at a low percent identity (60%) to remove sequences that were likely to be sequencing errors. Next, the closed reference OTU picking was applied on the filtered sequences. All sequences that did not match a reference at the closed reference step were then de novo clustered at 97% similarity. Singleton OTUs (i.e. sequences observed only once across the whole dataset) were discarded. Samples were rarefied to 3000 reads prior to diversity calculations; rarefaction was performed 10 times, and we report the mean Shannon diversity index for the 10 independent runs for each sample.

### Association of bacterial OTUs with symptoms of acute respiratory illness

We normalized the copy-number-corrected OTU read counts using cumulative sum scaling (CSS) (Paulson et al., 2013). Briefly, for each sample, the OTU counts were divided by the cumulative sum of counts up to the smallest percentile for which sample-specific count distributions were largely invariant (98.9^th^ percentile for our data). We then tested for differential abundance in ARI vs healthy samples, for each of 1,090 OTUs that were present in ≥10% of samples. A zero inflated Gaussian mixture model was fitted to the log transformed CSS-normalized OTU counts, separately for samples before and after 2 years of age (before 2 years: inclusive of samples at 2-year timepoint; after 2 years: from 2.5-year timepoint), using the R package *metagenomeSeq* (Paulson et al., 2013). We summarized the results for OTUs with an absolute fold change of >1.5 and FDR adjusted p-value <0.025 in either age strata. We picked eight representative OTUs to more precisely investigate how the associations changed over time, and modelled the longitudinal structure of the data using smoothing splines ANOVA with 100 permutations to assess significance (Paulson et al., 2017). All models for the ARI vs. healthy association were adjusted for age, season, gender, and any antibiotics within the last 4 weeks.

### Correlations between bacterial OTUs

We inferred correlation networks among the 1090 common OTUs using FastSpar (https://github.com/scwatts/fastspar), an efficient C++ implementation of the SparCC algorithm, which was designed to deal specifically with compositional data and produces more reliable and robust correlation estimates compared to Pearson or Spearman correlation especially in the case of low diversity samples (Friedman and Alm, 2012). SparCC uses a log ratio transformation and calculates correlations between OTUs in an iterative manner, under the assumption of a sparse network. Statistical significance of the correlation was assessed using 1000 bootstrap samples with exact p-value calculations based on the *permp* function in R package *statmod* (Phipson and Smyth, 2010). Correlation networks were generated across all samples, as well as separately for samples before and after 2 years of age, samples within each half yearly time periods, and samples with low abundance (<1%) of *Moraxella* OTU. We assessed differences in correlation before and after two years using the Fisher’s r-to-Z transformation (*cocor* R package).

### Within-individual dynamics

We first explored microbiome changes within each child in terms of transitions from a healthy sample to the next healthy sample half a year or a year later. Transitions which resulted in a MPG change indicated an abrupt shift in the major OTU (termed “unstable transitions”); stable otherwise. We also assessed the transitions using the Bray-Curtis dissimilarity calculated on the CSS-transformed OTU count matrix using the *vegan* R package (Oksanen et al., 2017), which represented subtle changes. Transition stability and distance were assessed for the effects of intervening ARI, LRI and antibiotics using GEE regression (logistic or linear where appropriate), adjusting for the age of first sample and difference in ages between samples.

We then investigated whether we could detect changes in the NPM prior to ARI symptoms, and how long these changes persisted after the illness. We grouped the healthy samples according to how soon after illness occurred (pre-illness: 1-2 weeks, 2-3 week, 3-4 weeks or >4 weeks) and how long after the last illness episode (post-illness: 1-2 months, 2-4 months, 4-6 months, 6-12 months, or >12 months). We used GEE logistic or linear regression to model (i) assignment to specific illness-associated MPGs or (ii) log abundance of specific illness-associated OTUs, against time to ARI/URI/LRI (separately for each pre-illness time category compared to all other healthy samples), and adjusted for time post-illness, gender, age, season, recent antibiotics, and any virus.

Lastly, we examined per child, if the proportion of illness-associated *Moraxella*, *Haemophilus* and *Streptococcus* MPGs in their healthy asymptomatic samples over different time periods (6 months to 2 years, and 2.5 years to 4 years) was associated with LRI frequency and subsequent wheeze phenotypes (wheeze at age 5 years or transient wheeze). Logistic regression was used to model (i) the proportion of illness-associated MPGs (as a binary: ≥50% versus <50%, excluding children with <2 healthy samples in each corresponding time period) against LRI and febrile LRI frequency in years 1, 2, 3 and 4, (ii) wheeze phenotypes against proportion of illness-associated MPGs (as quartiles), adjusting for LRI frequency. Separate models were fit for children with and without early allergic sensitization by 2 years of age.

### Virus detection

A second aliquot from 736 healthy, 583 URI and 789 LRI samples from the first three years (72%, 76% and 60% of all healthy, URI and LRI samples, respectively for which we had 16S profiles), were prepared for viral detection via reverse transcriptase polymerase chain reactions (PCR). Target organisms were: human rhinoviruses (RV); other picornaviruses (coxsackie, echo and enteroviruses); coronaviruses 229E and OC43; respiratory syncytial virus (RSV); influenza A and B; parainfluenzaviruses 1-3; adenoviruses and human metapneumovirus (HMPV). Primers, probes and PCR assay conditions have been previously described (Bochkov et al., 2014; Kusel et al., 2006; Kusel et al., 2007).

### Statistical Methods

All statistical analyses were performed using R (R Core Team, 2015) unless otherwise stated. Association analyses that involved multiple samples from the same subject were modelled using generalized estimating equations (GEE) with unstructured correlation and robust standard errors, where possible. In the case of non-convergence due to insufficient sample size, the ordinary logistic regression was used.

Potential confounders were included in the model. We used the Benjamini-Hochberg false discovery rate method (FDR) (Benjmini, 1995) or Bonferroni correction where multiple testing p-value adjustments were needed, as stated. All boxplots shown use the Tukey format, in which the bottom and top of the box represents the lower and upper quartiles respectively and the ends of whiskers represents the lowest/highest datum still within 1.5 interquartile range of the lower/upper quartile.

### Definition of variables used in statistical analyses

1. Wheeze at age 5: Presence of wheeze in the last 12 months recorded in the 5 year questionnaire.
2. Transient wheeze: Any wheeze in the first three years, but no wheeze in the 5^th^ year.
3. Early allergic sensitization: Any allergen-specific IgE levels > 0.35 kU/L by two years of age (at any of 6 month, 1 year or 2 years timepoints).
4. Season: According to month of collection: spring (September–November), summer (December–February), autumn (March–May) or winter (June–August).
5. Recent antibiotics: Any record of antibiotics intake within the last 4 weeks prior to sample collection.

## ACKNOWLEDGEMENTS

Supported in part by the Victorian Government’s Operational Infrastructure Support Program. This work was supported by the NHMRC of Australia (Project Grant #1049539 to MI and KEH, Fellowships #1061409 to KEH and #1061435 to MI).

## DATA AVAILABILITY

Sequencing data for this study, cleaned for human reads, has been deposited in the NCBI GenBank (accession TBA).

